# EFA6 regulates selective polarised transport and axon regeneration from the axon initial segment

**DOI:** 10.1101/150037

**Authors:** Richard Eva, Hiroaki Koseki, Venkateswarlu Kanamarlapudi, James W. Fawcett

## Abstract

It is not clear why central nervous system (CNS) axons lose their intrinsic ability to regenerate with maturity, whilst peripheral (PNS) axons do not. A key difference between these neuronal types is their ability to transport integrins into axons. Integrins can mediate PNS regeneration, but are excluded from adult CNS axons along with their rab11 positive carriers. We reasoned that this exclusion might contribute to the intrinsic inability of CNS neurons to regenerate, and investigated this hypothesis using laser axotomy. We identify a novel regulator of selective axon transport and regeneration, the ARF6 GEF EFA6. EFA6 exerts its effects from a previously unreported location within the axon initial segment (AIS). EFA6 does not localise here in DRG axons, and in these neurons, ARF activation is counteracted by an ARF-GAP which is absent from the CNS, ACAP1. Depleting EFA6 from cortical neurons permits endosomal integrin transport and enhances regeneration, whilst overexpressing EFA6 prevents DRG regeneration. Our results demonstrate that ARF6 is an intrinsic regulator of regenerative capacity, implicating EFA6 as a focal molecule linking the axon initial segment, signalling and transport.

**Summary Statement:** We identify a novel resident of the axon initial segment, EFA6. This functions to prevent growth-promoting molecules from entering mature CNS axons. Removing EFA6 elevates the axon’s regenerative potential.

## Introduction

Axons in the brain and spinal cord do not regenerate after injury because of a low intrinsic capacity for growth and extrinsic inhibitory factors (Fitch and Silver, 2008; Geoffroy and Zheng, 2014). Targeting inhibitory factors can promote recovery through sprouting and plasticity (Schwab and Strittmatter, 2014; Silver et al., 2014), but these interventions need to be combined with a strategy that promotes long range growth to optimise functional recovery. A number of studies have therefore focused on enhancing intrinsic regenerative capacity. These studies have identified signalling pathways and genetic factors that can be targeted to promote regeneration (Lindner et al., 2013; Liu et al., 2011; Moore and Goldberg, 2011), however regenerated axons often fail to reach their correct targets (Pernet and Schwab, 2014) and the cellular mechanisms downstream of these regeneration regulators are not completely understood. Investigating the mechanisms governing regenerative ability will help to explain how single interventions can orchestrate the numerous changes required to convert a dormant fibre into a dynamic structure capable of long range growth (Bradke et al., 2012).

Neurons in the peripheral nervous system (PNS) have a much greater capacity for regeneration, and can regenerate over long distances through the spinal cord if provided with an appropriate, activated integrin (Cheah et al., 2016). Integrins are adhesion molecules that mediate PNS regeneration, but are excluded from CNS axons after development (Andrews et al., 2016; Franssen et al., 2015). They are transported into PNS axons in recycling endosomes marked by the small GTPase rab11 (Eva et al., 2010). Rab11 governs the trafficking of many growth-promoting molecules (Solis et al., 2013; Wojnacki and Galli, 2016), and is necessary for growth cone function during development (Alther et al., 2016; van Bergeijk et al., 2015). However, rab11 is also excluded from mature CNS axons (Franssen et al., 2015; Sheehan et al., 1996). We reasoned that this restriction could be a major cause of regeneration failure, and that its reversal might be one of the mechanisms required to convert a quiescent axon into one capable of growth. Using a cell biology approach, we set out to determine how integrin and rab11 exclusion is controlled, whether it plays a part in governing regenerative ability, and how its regulation differs in PNS *vs.* CNS neurons.

Integrins are excluded from axons by two seemingly separate mechanisms involving ARF6 and the axon initial segment (AIS) (Franssen et al., 2015). We wondered whether these two mechanisms might be linked, and whether they might play a role in regulating regeneration. We identify the AIS-enriched ARF6 GEF EFA6 as a key molecule controlling selective axon transport and regenerative ability in CNS neurons. ARF6 and rab11 function as part of a complex, with ARF6 activation regulating transport direction (Montagnac et al., 2009). We used live cell imaging to determine that EFA6 regulates not only integrin transport, but also the transport of recycling endosomes marked by rab11. Removing EFA6 allows transport throughout CNS axons, and enables them to regenerate more efficiently after *in vitro* laser axotomy. This led us to understand why sensory axons have a much higher ability to regenerate than CNS neurons. In sensory neurons there is no transport block (Andrews et al., 2016). We show that EFA6 is not enriched in the initial part of PNS axons, but is instead present at low levels along the axon shaft. In these neurons, EFA6 activity is counteracted by an ARF6 inactivator (GAP) which is not present in CNS neurons, ACAP1. This is expressed at high levels throughout PNS axons. Overexpression of EFA6 inhibits PNS regeneration, as a result of its ARF GEF activity. Our results demonstrate that EFA6 and ARF6 are intrinsic regulators of regenerative capacity, and that they can be targeted to restore transport and promote regeneration.

## Results

### EFA6 localises to the axon initial segment, and activates axonal ARF

Our hypothesis was that the exclusion of integrins and rab11 positive recycling endosomes from adult CNS axons contributes to their inability to regenerate; that restoring the ability of CNS axons to transport growth-promoting machinery should boost their intrinsic regenerative ability. We aimed to identify novel targets for promoting axon transport and regeneration in CNS neurons. This would mean that in future, integrins might be used to promote guided regeneration of CNS axons through the spinal cord, as has already been achieved for sensory axons (Cheah et al. 2016). Finally, we anticipated that investigating CNS and PNS neurons might start to explain why these two different classes of neurons have different regenerative abilities, and identify mechanisms that may be functioning downstream of known regulators of regeneration.

We have previously found that several influences are responsible for excluding integrins from CNS axons (cortical neurons), including the axon initial segment (AIS) and a mechanism involving dynein dependent retrograde transport regulated by the small GTPase ARF6 (Franssen et al., 2015). An ARF6-rab11-JIP3/4 complex is known to control the direction of recycling endosome transport (Montagnac et al., 2009). We aimed to identify a single molecule which can be targeted to facilitate transport and promote regeneration, which might be functioning as a focal point to regulate signalling and transport mechanisms from the AIS. We reasoned that there may be an ARF6 activator in the AIS which prevents axonal integrin transport by stimulating retrograde transport, and unites these apparently unconnected mechanisms. For this work we have used a model of progressive regeneration failure in maturing cortical neurons validated in our previous research (Franssen et al. 2015, Koseki et al. 2017).

Of the known ARF6 guanine-nucleotide exchange factors (GEFs), EFA6 is strongly upregulated as neurons mature and develop selective polarised transport (Choi et al., 2006). It has a similar membrane targeting region to AIS spectrin (Derrien et al., 2002), and opposes axon regeneration in *c.elegans* (Chen et al., 2011). We used immunofluorescence to examine EFA6 localisation in cortical neurons differentiating *in vitro*. From 7 days *in vitro* (DIV) EFA6 was enriched in the initial part of the axon, where it colocalised with the AIS marker neurofascin (Fig. 1A). It was also present at lower levels throughout dendrites and the cell body, as previously reported (Choi et al., 2006) (Fig. 1A and B). EFA6 is an ARF6 GEF (Macia et al., 2001), which also regulates microtubules in *c. elegans* (Chen et al., 2011). We therefore investigated whether EFA6 was regulating ARF activation and/or microtubule dynamics. To investigate EFA6 GEF activity, we visualised activated ARF using the ARF binding domain (ABD) of GGA3 fused to a GST tag in 14DIV neurons. GGA3 is a coat protein that interacts only with the active form of ARF. ARF activation was not restricted to the AIS; instead we found a strong signal throughout axons (Fig. 2A). Importantly, this signal was not evident at 4DIV (when integrins and rab11 are transported into cortical axons). At this stage, when EFA6 was not enriched in the AIS, sparse vesicular structures were observed along the axons and these diminished at growth cones (Fig. 2B). Imaging at higher magnification confirmed that active ARF was detected uniformly along axons at the later developmental stage (Fig. 2C). To determine whether EFA6 was involved in axonal ARF activation in differentiated neurons, we depleted EFA6 with shRNA (Fig. S1). This led to a strong reduction in ARF activation throughout the axon, but did not affect total ARF6 (Fig. 2D). EFA6 preferentially activates ARF6 (Macia et al., 2001), so our finding suggests that EFA6 functions to activate ARF6 throughout axons, despite being restricted to the AIS. We next examined whether rodent EFA6 regulates microtubule dynamics by live imaging of the microtubule end-binding protein EB3-GFP, both in the AIS and throughout the axons of neurons expressing either control or EFA6 shRNA. We found that EB3-GFP was enriched in the AIS of control transfected neurons, as previously reported (Leterrier et al., 2011). Silencing EFA6 with shRNA had no effect on this distribution (Movies 1 and 2), suggesting that EFA6 does not affect microtubule stabilisation within the AIS (Leterrier et al., 2011). As a result of the high density of EB3 in the AIS, we did not detect comets here, even after depletion of EFA6. EB3-GFP comets were detected more distally into axons, but silencing EFA6 had no effect on either the number or behaviour of comets (Fig. S2). These data suggest that the developmental rise of EFA6 in the AIS leads to activation of ARF6 throughout mature CNS axons. However, rodent EFA6 does not regulate axonal microtubule dynamics as has been observed in *c. elegans*. This is consistent with the absence of the microtubule binding domain in mammalian EFA6.

**Fig. 1.**
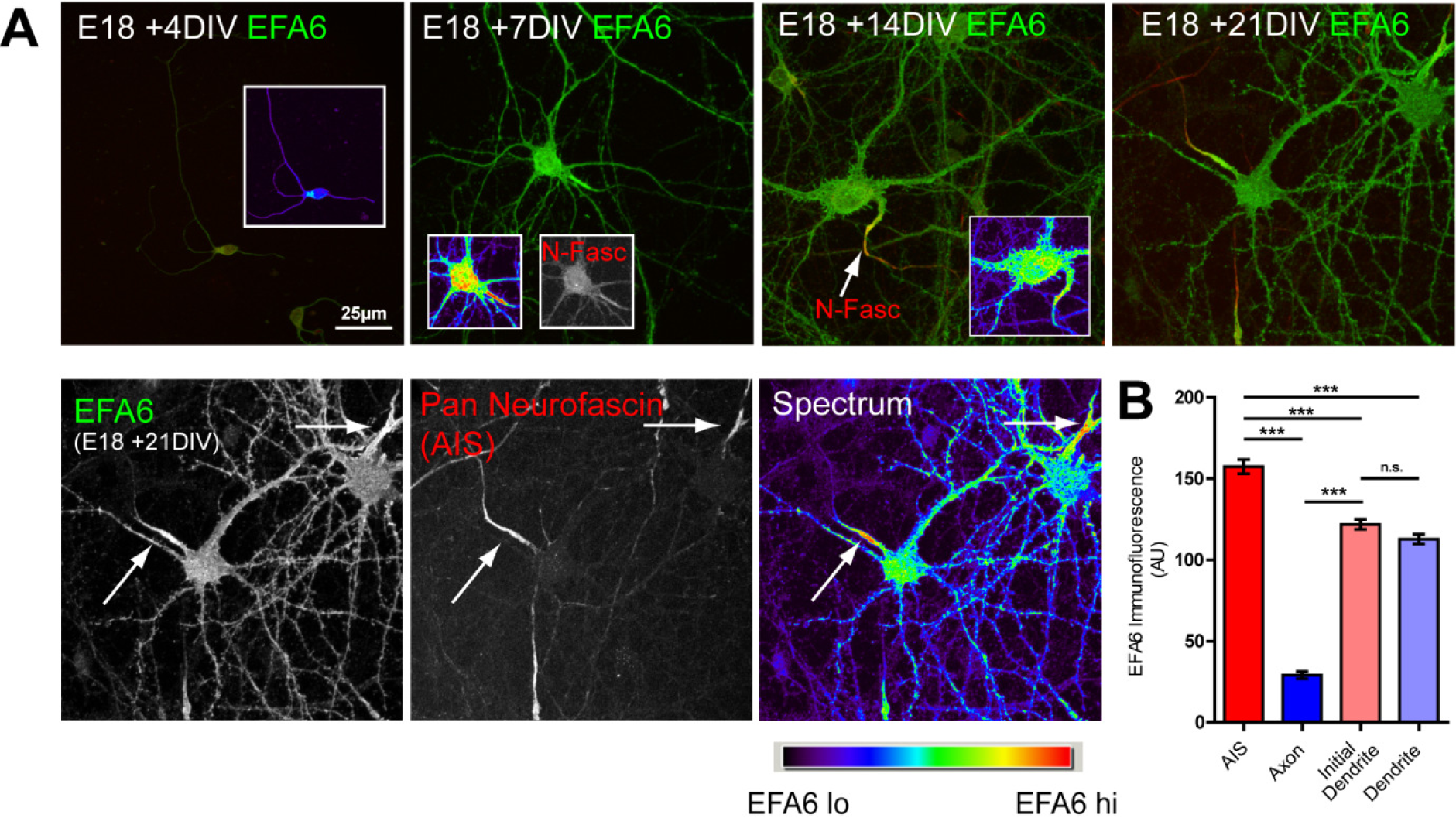
EFA6 is enriched in the axon initial segment. **(A)** Immunolabelling of EFA6 and neurofascin in cortical neurons (E18 +4 to 21DIV). EFA6 is expressed at low levels throughout the cell at an early stage (E18 +DIV4). Expression increases with maturity, and EFA6 is enriched at the axon initial segment from +7 DIV onwards. EFA6 green colour, pan-neurofascin red colour (to identify the axon initial segment). Arrows indicate the AIS. Spectrum colouring indicates highest signal in the AIS. **(B)** Quantification of fluorescence intensity of EFA6 in the AIS, axon, initial dendrite and dendrite. n=45 neurons from three experiments. *** p<0.0001, using Anova and Bonferroni**’**s comparison test.

**Fig. 2.**
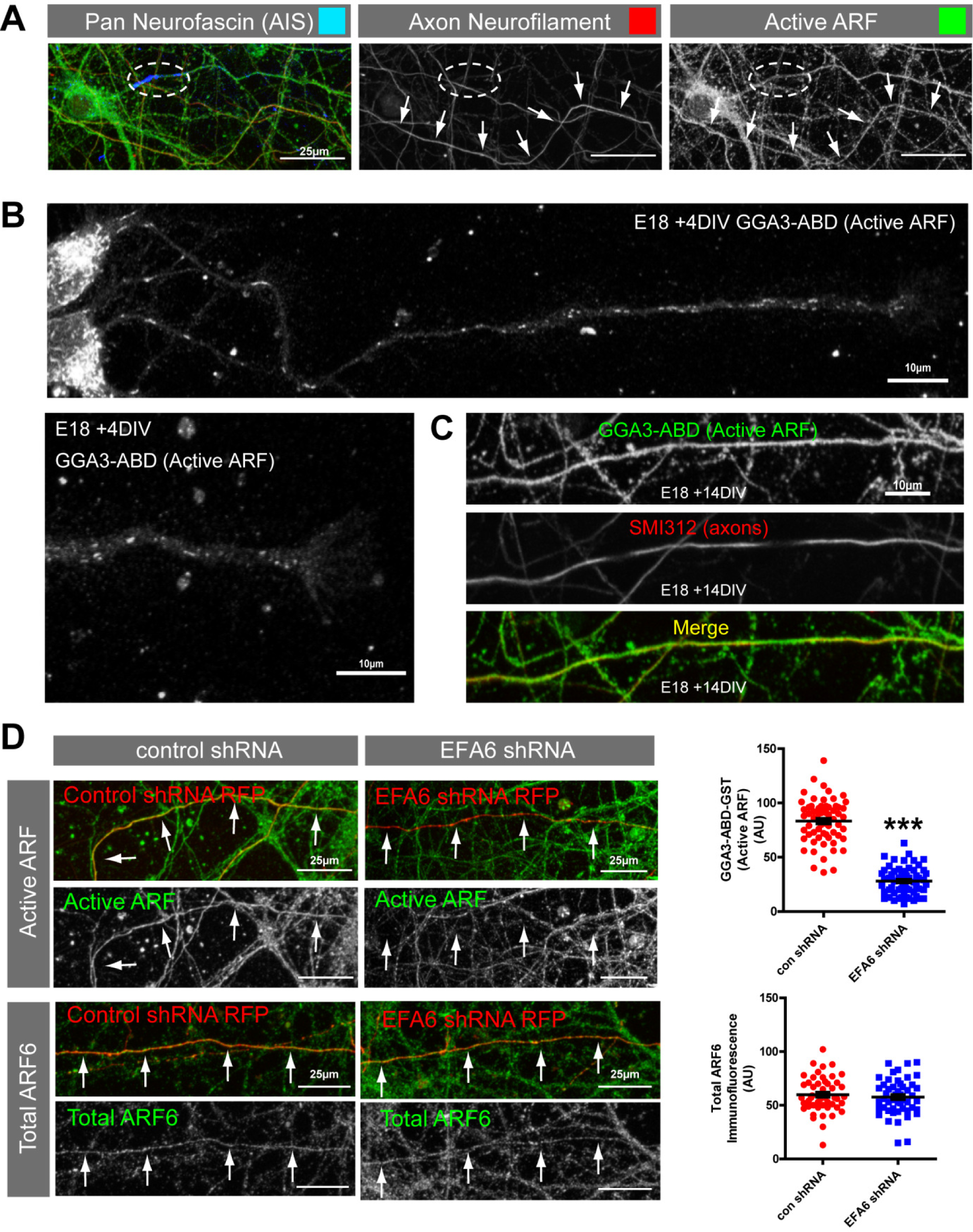
EFA6 activates ARF throughout mature CNS axons. **(A)** Active ARF (GGA3-ABD-GST, green) in proximal axons (neurofascin in blue), and distal axons (axon neurofilaments, red). Ellipse indicates the initial segment, arrows indicate a distal axon. **(B)** In young neurons (+4DIV), active ARF is detected in sparse tubulo-vesicular structures throughout developing axons which diminish at the growth cone. **(C)** In differentiated neurons (+14DIV) active ARF (GGA3-ABD-GST + anti-GST) is distributed uniformly throughout axons (as indicated by immunolabelling with SMI312 for axonal neurofilaments). **(D)** Active ARF and total ARF6 in axons of neurons expressing either control shRNA or shRNA targeting EFA6 (red). Active ARF or ARF6 is shown in green, as indicated. Arrows indicate axons. Quantification of axonal ARF activity and total ARF6 in neurons expressing either control or EFA6 shRNA. n=61 to 64 neurons (active ARF), *** p<0.0001, students t-test. n=50 for total ARF6 quantification.

### EFA6 directs integrins away from axons

Whilst endogenous integrins are restricted to dendrites in differentiated CNS neurons, overexpressed integrins enter proximal axons where they exhibit predominantly retrograde transport. Retrograde transport is one mechanism through which a polarised distribution can be achieved in neurons (Guo et al., 2016; Kuijpers et al., 2016). The direction of axonal integrin transport is regulated by ARF6 activation state, such that elevated ARF6 activation causes retrograde transport (Franssen et al., 2015). Removal of the chief ARF6 activator, EFA6, should therefore facilitate anterograde integrin transport. We used live spinning disc confocal microscopy to image and analyse axonal integrin movement at three points (AIS, proximal and distal) using α9 integrin-GFP (this integrin promotes spinal sensory regeneration (Cheah et al., 2016)), in the presence of EFA6-shRNA or control. Anterograde transport was almost undetectable in control transfected neurons. These exhibited predominantly retrograde and static vesicles, and there was a rapid decline in integrin levels with distance (Fig. 3A and B, Fig. S3, Movies 3 and 4). Depleting EFA6 initiated anterograde transport, diminished retrograde transport, and increased integrins in all segments of the axon (average 8.3 vesicles per section/AIS, 7.9/proximal, 7.1/distal). Endogenous β1 integrin (the binding partner of α9) also entered axons. Measurement of the axon-dendrite ratio showed that depleting EFA6 lead to integrins being present in axons at a similar level to dendrites (axon-dendrite ratio changing from 0.24 in control transfected neurons, to 0.95 in neurons expressing EFA6 shRNA) (Fig. 3C and D). Removing the ARF6 GEF EFA6 therefore enables anterograde integrin transport and increases integrin levels throughout the axon.

**Fig. 3.**
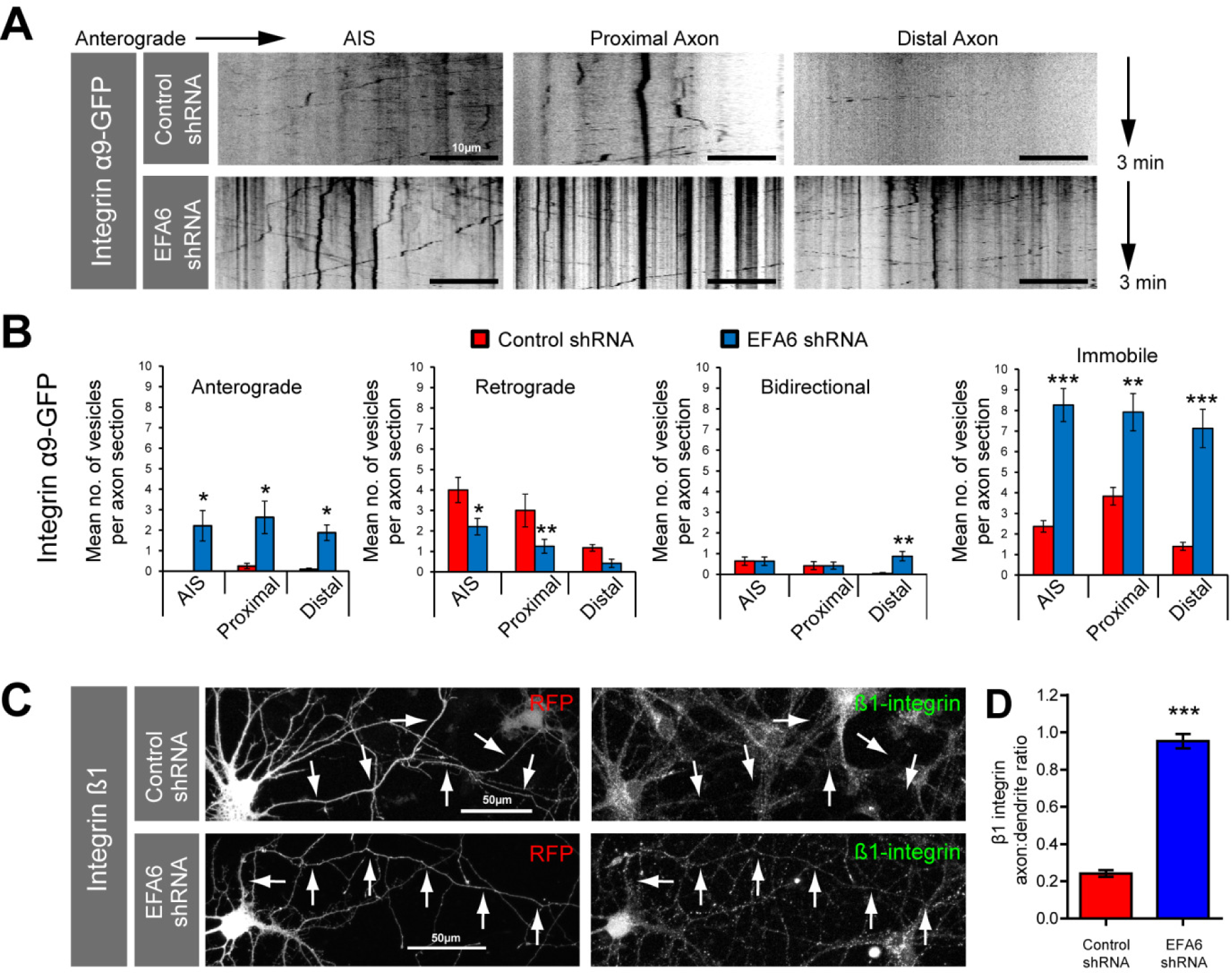
Depletion of EFA6 promotes axon transport of α9 and β1 integrins. **(A)** Kymographs showing dynamics of α9 integrin-GFP in the AIS, proximal and distal axon of neurons expressing control or EFA6 shRNA. **(B)** Quantification of α9 integrin-GFP axon transport n=12 to24 neurons per condition, total 1024 vesicles. ***, **, * indicates p<0.001, p<0.01, p<0.05 respectively, using Anova and Bonferroni’s comparison test. **(C)** Immunolabelling of β1 integrin in neurons expressing control or EFA6 shRNA. Arrows indicate axons. **(D)** Quantification of axon-dendrite ratio of endogenous β1 integrin after EFA6 silencing. n=71 neurons from three experiments. p<0.0001 by students t-test.

### EFA6 directs rab11 endosomes away from axons, but does not affect APP transport

Axonal integrins traffic via recycling endosomes marked by rab11. This small GTPase is necessary for growth cone function during developmental axon growth in the CNS and involved in axon regeneration (Eva et al., 2010; Nguyen et al., 2016; van Bergeijk et al., 2015). However, it is excluded from mature CNS axons (Franssen et al., 2015; Sheehan et al., 1996). Rab11 and ARF6 cooperate to control microtubule-based transport direction (Montagnac et al., 2009) and axon growth (Eva et al., 2012; Eva et al., 2010; Suzuki et al., 2010). From its effects on integrin transport, we reasoned that EFA6 may be pivotal in directing rab11 away from axons. As with integrins, some overexpressed rab11-GFP leaks into CNS axons (detectable as vesicular punctae) although at much lower levels than dendrites. This allows for analysis of axonal vesicle dynamics (Fig. 4A). In the AIS individual vesicles could not be distinguished (Fig. 4A, AIS kymograph). Beyond the AIS, anterograde transport was minimal in control transfected cells, with the majority of vesicles moving either retrograde, bidirectional, or remaining immobile. Overall transport declined with distance (Average 12.3 vesicles per proximal section, 6.7 per distal section, Fig. 4B). Expressing EFA6-shRNA stimulated anterograde transport. There was also a reduction in retrograde transport in the proximal axon, an increase in bidirectional movement in the distal axon, and an increase in immobile punctae throughout the axon (Fig. 5A and B, and extended data figure 4 and 5-1). These transport changes permitted endogenous rab11 to enter axons, altering the axon:dendrite ratio from 0.35 to 0.94 (control vs. EFA6 shRNA) (Fig. 4C and D). EFA6 therefore functions to limit the axonal localisation of integrins and rab11 positive endosomes. To confirm the selectivity of these effects we also analysed the transport of amyloid precursor protein (APP), a molecule which targets to CNS axons (Chiba et al., 2014) that is not normally found in rab11 endosomes (Steuble et al., 2012). We found no differences in APP axon transport dynamics between neurons expressing control or EFA6 shRNA (Fig. 4E and F), indicating that EFA6 depletion does not alter global axon transport.

**Fig. 4.**
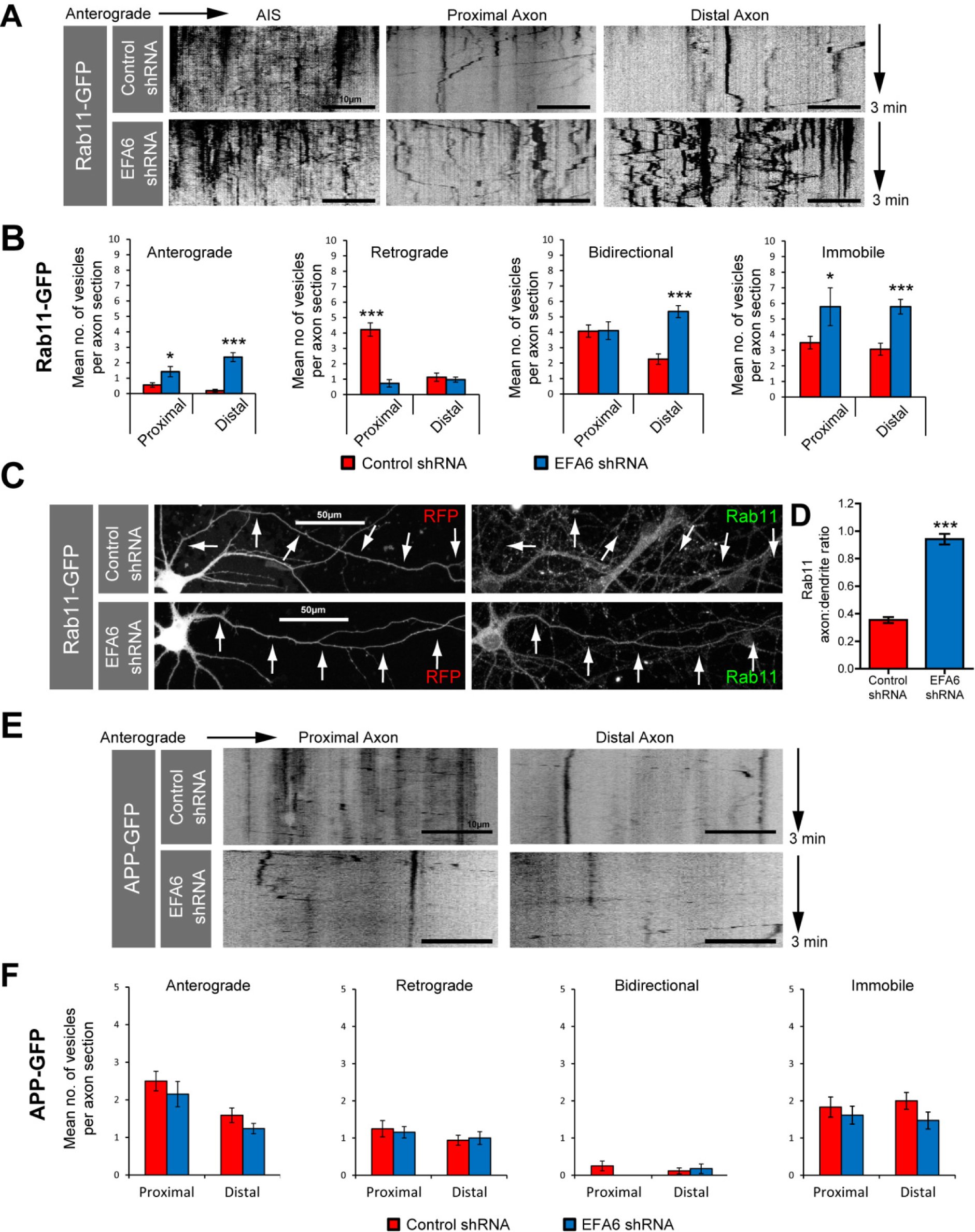
Depletion of EFA6 promotes axon transport of rab11, and not APP. **(A)** Kymographs showing dynamics of rab11-GFP in the AIS, proximal and distal axon of neurons expressing control or EFA6 shRNA. **(B)** Quantification of rab11-GFP axon transport, n=19 to27 neurons per condition. *** p<0.001, * P<0.05 respectively, using Anova and Bonferroni’s comparison test. **(C)** Immunolabelling of rab11 in neurons expressing either control or EFA6 shRNA. Arrows indicate axons. **(D)** Quantification of endogenous rab11 axon-dendrite ratio after EFA6 silencing. n=71 neurons from three experiments. p<0.0001 by students t-test. Also see associated Figure SI2. **(E)** Kymographs showing dynamics of APP-GFP in the proximal and distal axons of neurons expressing either control shRNA or shRNA targeting EFA6. **(F)** Quantification of APP-GFP vesicle movements in the proximal and distal axon. No statistical difference was found between neurons expressing control- or EFA6-shRNA using ANOVA and Bonferroni’s comparison test.

### EFA6 depletion enhances regeneration of CNS axons

Rab11, integrins and reduced ARF6 activity are all beneficial for axon growth (Cheah et al., 2016; Eva et al., 2012; Eva et al., 2010; Franssen et al., 2015; Gardiner, 2011; van Bergeijk et al., 2015). We therefore asked whether silencing EFA6 would intrinsically enhance regeneration. *In vitro* single-cell axotomy enables detailed study of intrinsic regenerative capacity, allowing morphological evaluation of the regenerative response after injury (Gomis-Ruth et al., 2014). We developed an *in vitro* laser axotomy protocol for analysing the regeneration of individually axotomised cortical neurons (Koseki et al., 2017). E18 rat cortical neurons were cultured on glass imaging dishes, transfected at 10DIV, and used for experiments at 14-17DIV. We used this system to study the effects of EFA6 depletion on regeneration of cortical neurons after laser axotomy (Fig. 5A and S4). When regeneration was successful, we recorded growth cone size, time taken to regenerate, and distance grown after regeneration (within the experimental time frame). We recorded whether axons formed stumps or motile end bulbs after injury when regeneration failed (Fig. 5B-C). By 14DIV integrins and rab11 are mostly excluded from axons, and their ability to regenerate is limited (Koseki et al., 2017). Neurons expressing EFA6-shRNA showed a substantial increase in regeneration with 57.6% of neurons regenerating their axons within 14 hours, compared to 27.4% of neurons expressing control-shRNA (Fig. 5D-F, Movies 5 and 6). They also developed larger growth cones, extended their axons over greater distances and initiated regeneration more rapidly than cells expressing control-shRNA (Fig. 5D, E, H-J). EFA6 shRNA treated neurons tended to form motile bulbs when they failed to regenerate (84.8%), whereas control transfected cells tended to from immobile stumps (55%) (Fig. 5B, C, D, E and G). Depleting EFA6 therefore raises the regenerative capacity of differentiated cortical neurons.

**Fig. 5.**
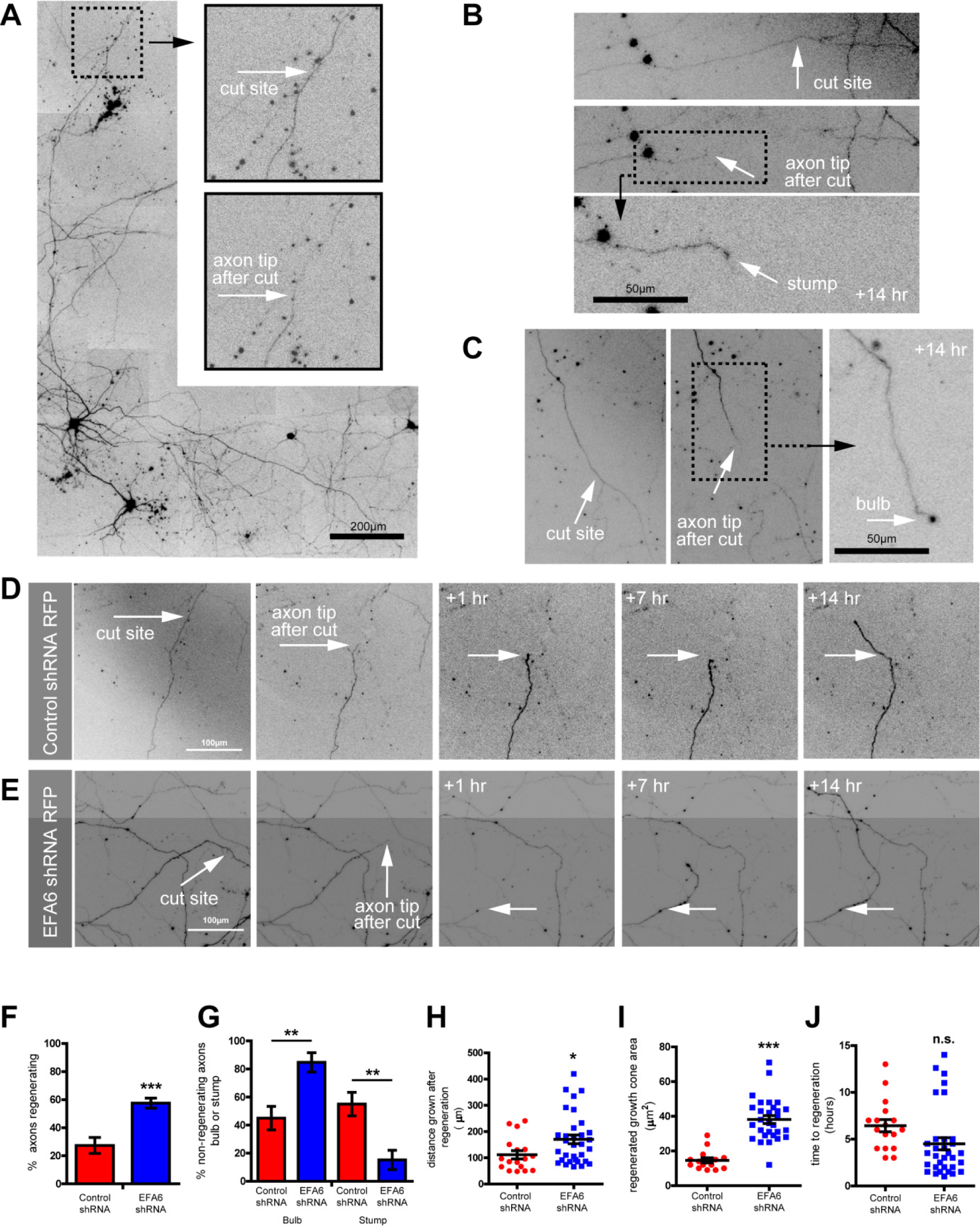
Depleting EFA6 promotes axon regeneration in CNS neurons. **(A)** Example of a neuron used for CNS axotomy experiments, indicating the site chosen for laser ablation (typically >1000μm distal, on an unbranched section of axon). Fluorescent signal is control shRNA-RFP. **(B)** Example of regeneration failure, and formation of stump. **(C)** Example of post-axotomy end bulb formed after axotomy. **(D)** Neuron expressing control shRNA, showing axotomy followed by regeneration. Note the small growth cone (typically <20μm^2^ and regeneration of <100μm in 14hrs. **(E)** Neuron expressing EFA6 shRNA, showing axotomy followed by regeneration >100μm in 14hrs, with growth cones typically >40μm^2^. (F-J) Quantification of regenerative response of cut axons of neurons expressing either control or EFA6 shRNA. **(F)** Percentage of axons regenerating within a 14hr period. p<0.001 by fishers exact test. n=59 to 63 neurons. **(G)** Percentage of failed axons bulb vs. stump, p<0.01 by fishers exact test. **(H)** Distance grown after regeneration, p<0.05 by t-test. **(I)** Area of regenerated growth cones, p<0.0001. **(J)** Time taken to establish a growth cone and regenerate >50μm.

### ARF6 is an intrinsic regulator of regenerative capacity

As EFA6 contributes to the low regenerative capacity of CNS neurons, we reasoned that neurons that can regenerate their axons should either have less axonal EFA6, or a means of counteracting its effects. In the PNS, adult dorsal root ganglion (DRG) neurons have regenerative axons which permit integrin and rab11 transport (Andrews et al., 2016; Gardiner, 2011). When we examined EFA6 in adult DRG neurons we found remarkably high levels in the cell body, and lower levels throughout axons (Fig. 6A). We speculated that this may be counteracted by an ARF6 inactivator and investigated ACAP1, a known regulator of integrin traffic which we previously used to manipulate integrin transport in DRG axons (Eva et al., 2012; Li et al., 2005). ACAP1 was present in adult DRG neurons, throughout axon shafts and at growth cones, but was absent from cortical neurons (Fig. 6A). This led us to compare ARF activation in DRG and differentiated cortical neurons. We found that ARF activation was evident in DRG axons, but at a lower level than cortical axons (Fig. 6B). This suggests that PNS axons may be better regenerators due to expression of an ARF6 inactivator. This hypothesis predicts that elevating ARF6 activation in DRG axons would inhibit regeneration. We used laser axotomy to injure the axons of adult DRG neurons *in vitro*. We examined regeneration in the presence of overexpressed GFP, EFA6-GFP, or EFA6 E242K-GFP (ARF6 activation incompetent) (Fig. 7). Control DRG axons regenerate rapidly, so that by 2hrs after injury 68.6% of GFP expressing axons had developed new growth cones (Fig. 7A, C and E, Movie 7). Overexpression of EFA6 led to a dramatic reduction in regenerative capacity with only 19.2% of axons regenerating growth cones (Fig. 7B, D and E, Movie 8). This effect was primarily due to EFA6 GEF activity, as expression of EFA6 E242K did not have the same effect, allowing 50.5% of axons to regenerate (Fig. 7E and S5). This suggests that EFA6 opposes regeneration principally by virtue of its GEF activity towards ARF6. The data demonstrate that ARF6 activation state plays a central role in regulating the regenerative capacity of DRG neurons. Taken together with our findings in differentiated cortical neurons, our data suggest that the activation state of ARF6 is an intrinsic regulator of axon regeneration, responsible for the exclusion of Rab11 vesicles and their contents from CNS axons.

**Fig. 6.**
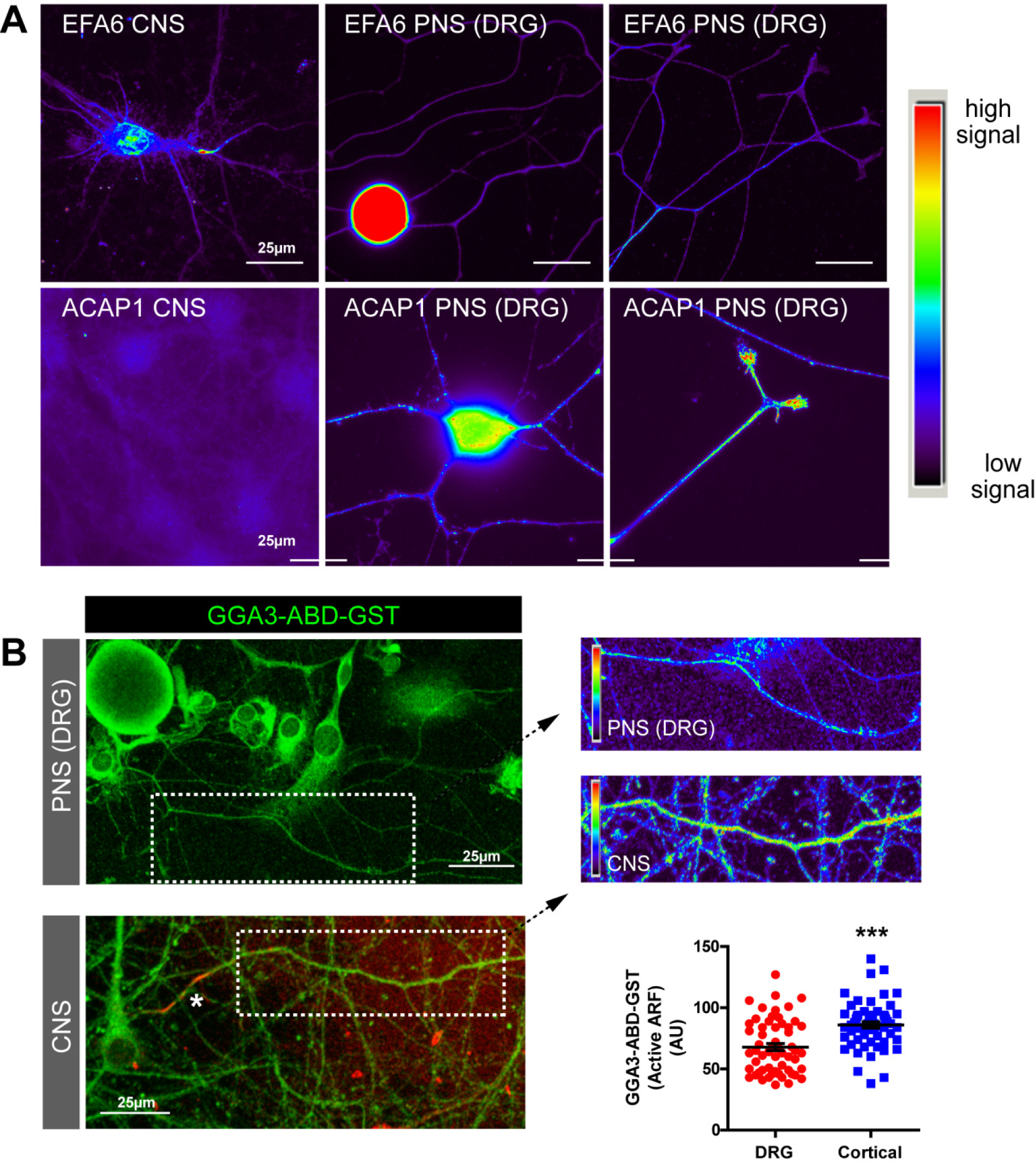
ARF6 is regulated differently in CNS *vs*. PNS neurons. **(A)** Cortical neurons and adult DRG neurons immunolabelled for EFA6 (upper panels) or ACAP1 (lower panels). Both neuronal types were labelled and imaged identically to allow comparison of fluoresent signal. Images represent two independent immunolabelling experiments. **(B)** Axons of DRG and cortical neurons (+DIV10) labelled with GGA3-ABD-GST to detect active ARF. Graph shows quantification of ARF activation in the two axon types. n=58(DRG) and 60(cortical). ***p<0.001 by T-test.

**Fig. 7.**
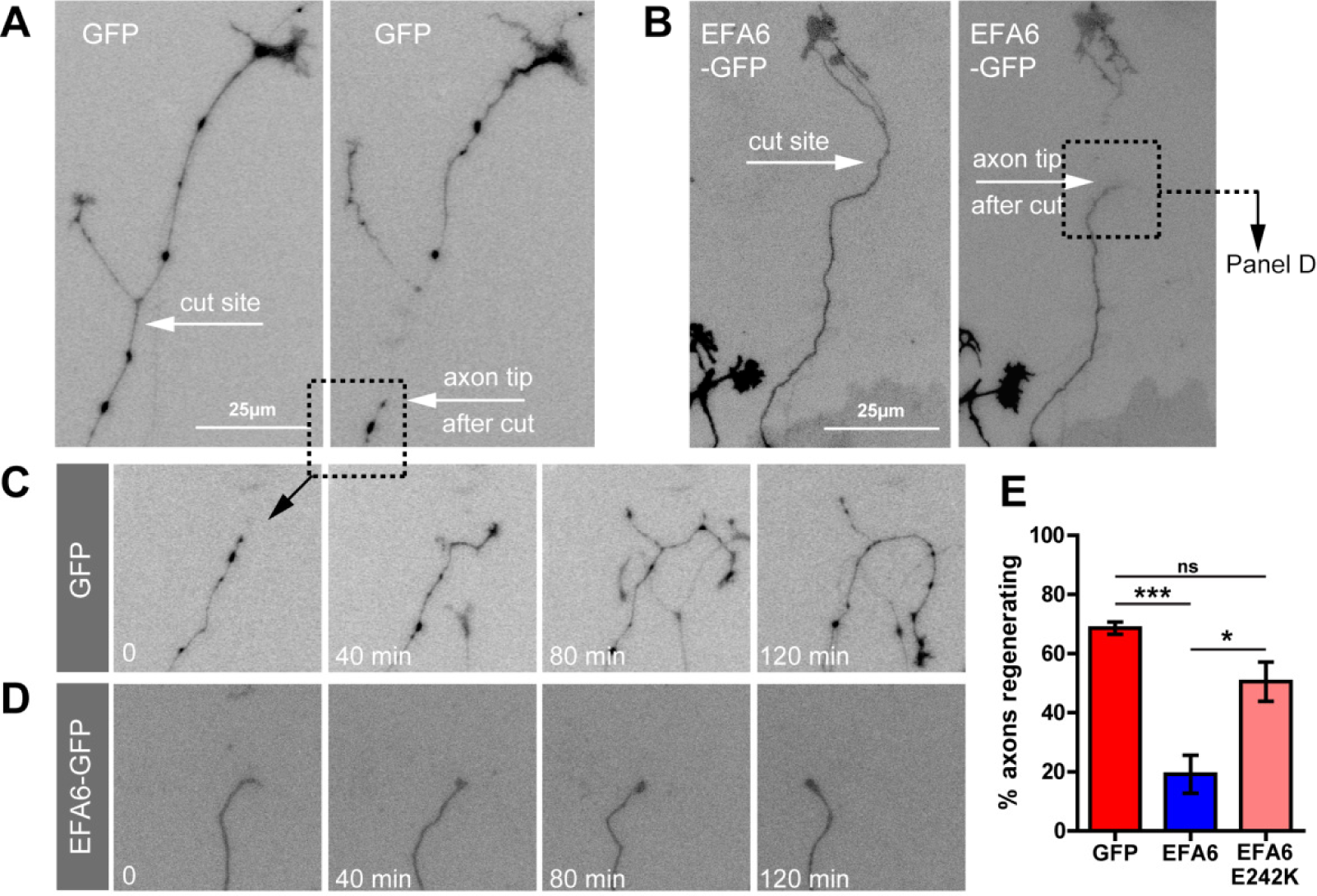
Elevated ARF6 activation inhibits regeneration of adult DRG axons. **(A)** Cut DRG axon expressing GFP. **(B)** Cut DRG axon expressing EFA6-GFP. **(C)** Axon from panel A showing regeneration. **(D)** Axon from panel B showing failure to regenerate. **(E)** Quantification of axon regeneration of DRG neurons expressing either GFP (n=48), EFA6-GFP (n=44) or EFA6 E242K (EFA6 lacking the ability to activate ARF6) (n=31). ***p<0.0001, *p<0.05 by Fisher’s exact test.

## Discussion

Our data demonstrate that the exclusion of integrins and recycling endosomes from mature CNS axons plays an important role in limiting regenerative potential. We show that EFA6 is developmentally up-regulated and enriched in the AIS at a time when integrin transport becomes predominantly retrograde (Franssen et al., 2015), and neurons lose their ability to regenerate. From its site in the initial part of the axon, EFA6 functions to activate ARF6 throughout mature axons, leading to retrograde removal of integrins and rab11 endosomes. Removing EFA6 restores transport, and facilitates regeneration. These phenomena are specific to CNS neurons, as EFA6 is not enriched in the initial part of regenerative PNS axons (of sensory DRG neurons). Sensory neurons regulate axonal ARF6 differently, by expressing an ARF GAP which is absent from cortical neurons, ACAP1. Overexpressing EFA6 opposes regeneration in these neurons, principally by virtue of its GEF domain. Our findings start to explain, at a cellular level, why PNS neurons have a better capacity for regeneration than their CNS counterparts. The results implicate ARF6 as an intrinsic regulator of regenerative potential, and identify EFA6 as a novel target for promoting CNS axon regeneration.

### EFA6 activates axonal ARF to control selective polarised transport

We have found that EFA6 activates ARF6 throughout the axon despite being enriched in the AIS. How does EFA6 achieve ARF activation over long distances? This may involve a complex interaction with an additional ARF regulator, ARNO, which localises throughout the axon (Franssen et al., 2015). EFA6 is known to control a negative-positive feedback circuit between EFA6, ARF6 and ARNO. EFA6 is necessary to establish initial ARF activation, which is consequently maintained by ARNO (Padovani et al., 2014). In axons, ARF activation event may be spatially regulated, with initial activation occurring within the AIS, and subsequent activation maintained throughout the axon by ARNO. Axonal ARF activation may be necessary to aid neurotransmission, because ARF6 activation drives synaptic vesicles towards recycling rather than endosomal sorting, enabling maintenance of the readily releasable synaptic vesicle pool (Tagliatti et al., 2016).

The AIS is primarily responsible for initiation of the action potential, but is also involved in the polarised delivery of membrane proteins, ensuring the correct distribution of axonal and dendritic machinery as neurons mature (Bentley and Banker, 2016; Rasband, 2010). The molecular mechanisms through which this is achieved are not completely understood, but are reported to involve the actin and microtubule cytoskeleton (Arnold, 2009; Britt et al., 2016) and dynein dependent retrograde transport (Kuijpers et al., 2016). We have previously found that integrins are removed from axons by dynein dependent retrograde transport, and that lowering ARF activation reduces retrograde removal of integrins and allows modest anterograde transport. We also found that removing the AIS by silencing is central organiser, ankyrin G, also permits some anterograde transport, but we did not understood how these phenomena might be linked. Here we establish a mechanism for the removal of both integrins and rab11 endosomes, controlled by EFA6 from a location in the AIS. EFA6 is probably localised here by virtue of its membrane targeting motif, which has a high degree of homology with the membrane binding region of the AIS component, βIV spectrin (Derrien et al., 2002). Our current data suggest a novel model for selective distribution. The selectivity comes from the involvement of the small GTPases ARF6 and rab11, and likely involves the adaptor molecules JIP3 and 4. ARF6 forms a complex with rab11, and the adaptor molecules JIP 3 and 4: the activation state of ARF6 determines whether this complex associates with dynein or kinesin, and therefore its direction of transport (Montagnac et al., 2009). The JIP adaptors are part of a family of 4 proteins (JIPs 1-4,) which link cargo to motor proteins. JIPs 1 and 2 are similar to each other, but differ from JIPs 3 and 4 (Koushika, 2008). ARF6 is known to interact with JIP3 in cortical axons (Suzuki et al., 2010). Crucially ARF6 does not interact with JIPs 1 and 2 (Koushika, 2008), meaning that its activation will not affect the transport of cargo which interacts specifically with these two adapters. We have shown here that APP transport is not affected by EFA6 silencing. It is important to note that APP interacts with JIPs 1 and 2, and not JIP 3 or 4 (Chiba et al., 2014; Edwards et al., 2013). APP also traffics independently from rab11 (Chiba et al., 2014). We speculate that for the axonal transport of a molecule to be affected by ARF6, it needs to traffic through a rab11/ARF6 compartment, and also interact with JIP 3/4. The ARF-dependent control of entry to a specific cellular compartment is not without precedent. A similar mechanism regulates the entry of rhodopsin into primary cilia. In this case, a specific ARF inactivator (ASAP1) is required to permit ARF4 and rab11 dependent transport (Wang et al., 2012).

### Rab11, ARF6 and axon regeneration

Much is known about the mechanisms required for growth cone formation and subsequent axon growth, but is not understood why these mechanisms are not recapitulated after injury in the brain or spinal cord (Bradke et al., 2012). Our study demonstrates that the supply of growth promoting material in recycling endosomes is an important factor governing regenerative potential. An axon cannot rebuild a functional growth cone without the appropriate materials. It is well established that integrins are important for axon growth during development as well as for regeneration after injury in the PNS (Gardiner, 2011; Myers et al., 2011), and it would appear that the same can be said for rab11 positive recycling endosomes. These are necessary for growth cone function in CNS neurons during development (Alther et al., 2016; van Bergeijk et al., 2015), and now appear to participate in regeneration of CNS neurons axotomised *in vitro*. Some of rab11’s functions during axon growth may be due to the supply of integrins, however other growth-promoting molecules also traffic via rab11, including neurotrophin receptors, (Ascano et al., 2009; Lazo et al., 2013), and the pro-regenerative flotillin/reggie proteins (Bodrikov et al., 2017).

Our data imply a central role for ARF6 and EFA6 in determining the intrinsic regenerative ability of neurons. As CNS neurons mature, EFA6 is enriched in the initial part of the axon, leading to axonal ARF activation and retrograde removal of molecules necessary for growth. Conversely, adult PNS neurons regenerate better, have low levels of EFA6 in their axons, express an ARF6 inactivator, which is not found in cortical neurons (ACAP1), and permit rab11 and integrin transport (Eva et al., 2010). Overexpressing EFA6 in PNS neurons leads to a reduction in regenerative capacity (and increased retrograde transport, (Eva et al., 2012)), while decreasing EFA6 in cortical neurons restores regeneration. These combined results suggest that ARF6 functions as an intrinsic regulator of regenerative capacity, governed by its activation state. This novel finding is in keeping with a known intrinsic regulator of regenerative potential, the tumour suppressor PTEN. Deleting PTEN enhances regeneration in the CNS, partly through the PI3 kinase/AKT/mTOR pathway (Park et al., 2008). PTEN and PI3 kinase counteract each other to regulate the amounts of phosphatidylinositol phosphates (PIPs) PIP2 and PIP3. The majority of ARF6 GEFs and GAPs are regulated downstream of PIP2 or PIP3 (Randazzo et al., 2001), and the activity of EFA6 is strongly elevated in the presence of PIP2 (Macia et al., 2008). It is therefore possible that deletion of PTEN could result in less PIP2, and lowered EFA6 activity. We hypothesise that the expression profile of axonal ARF regulators and the phosphoinositide environment are crucial factors that control the axonal entry of regenerative machinery and its subsequent insertion onto the surface membrane. These are crucial factors that determine whether a damaged axon can reconstruct a functional growth cone to drive guided axon regeneration after injury.

## Materials and Methods

### Neuron cultures and transfection

Primary cortical neuron cultures were prepared from embryonic day 18 (E18) Sprague Dawley rats. Neurons were dissociated with papain (Worthington) for 8 min at 37°C, washed with HBSS and cultured in NeuralQ^®^ Basal Medium (AMSBio) supplemented with GS21 (AMSBio), and glutamax (Thermo). Cells were plated on glass-bottom dishes (Greiner) coated with poly-D-lysine. Culture dishes were incubated in humidified chambers to prevent evaporation of culture medium, allowing long-term culture (up to 28days in vitro (DIV)). DRG neuronal cultures were obtained from adult male Sprague Dawley rats. DRGs were incubated with 0.1% collagenase in DMEM for 90 min at 37°C followed by 10 min in trypsin at 37°C. DRGs were dissociated by trituration in a blunted glass pipette. Dissociated cells were then centrifuged through a layer of 15% bovine serum albumin (BSA), washed in DMEM, and cultured on 1 μg/ml laminin on poly-D-lysine coated glass-bottom dishes (Greiner) in DMEM supplemented with 10% fetal calf serum (Thermo), 1% penicillin/streptomycin, and 50 ng/ml nerve growth factor (NGF). Cortical neurons were transfected by oscillating nano-magnetic transfection (magnefect nano system, nanoTherics, Stoke-On-Trent, UK) as previously described (Franssen et al., 2015). For EFA6 silencing, cells were transfected at 10DIV, and experiments (imaging or axotomy) were performed between 14 and 17 DIV. Transfections of dissociated adult DRG neurons were performed *in situ* at 1DIV as previously described (Eva et al., 2012) using a Cellaxess in-dish electroporator (Cellectricon).

### DNA and shRNA constructs

Integrin alpha9 EGFP-N3 was obtained from Addgene (Addgene plasmid # 13600), deposited by Prof. Dean Sheppard (University of California, San Francisco, CA), previously characterised (Eva et al., 2012; Eva et al., 2010; Franssen et al., 2015). cDNA encoding human Rab11a was amplified by PCR introducing HindIII and BamHI restriction sites, and cloned into EGFP-C2 (Eva et al., 2010). APP-GFP was a gift from Prof. Michael Coleman (Cambridge University, Centre for Brain Repair)(Hung and Coleman, 2016). EB3-GFP was a gift from Prof. Casper Hoogenraad (Utrecht University, Netherlands) (Stepanova et al., 2003). Human EFA6 ORF in pFLAG-CMV6-EFA6 (Brown et al., 2001) was a gift from Dr. Julie Donaldson (Bethesda, MD). Human EFA6 cDNA was obtained by PCR amplification using pFLAG-CMV6-EFA6 as a template and was digested with EcoRI and SalI and cloned into the same sites of pEGFP-C1 (Clontech) to obtain pEGFPC1-EFA6 (Eva et al., 2012). Rat EFA6 (NM_134370) in pcDNA3.1-C-(k)DYK (FLAG tag) was obtained from Genscript. EFA6 E242K-GFP (Luton et al., 2004) was a gift from Frederic Luton (Valbonne, France). EFA6 silencing was achieved using shRNA targeting EFA6 in pRFP-C-RS (PSD gene NM_134370, targeting sequence cagtcctggattactcgcatcaatgtggt) (Origene). Non-effective 29- mer scrambled shRNA cassette in pRFP-C-RS vector (Origene) was used as a control.

### Antibodies

Guinea pig polyclonal EFA6A (1626) antibody was a gift from Prof. Eunjoon Kim (Daejeon, South Korea) previously characterised (Choi et al., 2006) and used in independent studies (Raemaekers et al., 2012; Sannerud et al., 2011). Other primary antibodies: mouse anti-neurofascin clone A12/18, NeuroMab (RRID:AB_10671311). Rabbit anti-Rab11 71-5300, Thermo. Rabbit anti-ARF6 ab77581 (Abcam,) Rabbit anti-GST ab19256 (Abcam,) mouse anti-FLAG ab18230 (Abcam). Mouse anti beta actin ab8226 (Abcam), rabbit anti-tRFP (Evrogen), anti-integrin ß1 clone EP1041Y (04-1109, Millipore), anti- pan-axonal neurofilaments mouse monoclocal SMI312 (Abcam ab24574). ACAP1 detected with goat anti Centaurin β1 ab15903 (Abcam). Secondary antibodies were alexa conjugates from Thermo used at 1:800. Secondary antibodies for western blotting were HRP conjugates from GE Life Sciences.

### Microscopy

Laser scanning confocal microscopy was performed using a Leica DMI4000B microscope, with laser scanning and detection achieved by a Leica TCS SPE confocal system controlled with Leica LAS AF software. Fluorescent and widefield microscopy was performed using a Leica DM6000B with a Leica DFC350 FX CCD camera and a Leica AF7000 with a Hamamatsu EM CCD C9100 camera and Leica LAS AF software. Leica AF7000 was also used for imaging of axon and growth cone regeneration after axotomy. Live confocal imaging was performed with an Olympus IX70 microscope using a Hamamatsu ORCA-ER CCD camera and a PerkinElmer UltraVIEW scanner for spinning disk confocal microscopy, controlled with MetaMorph software.

### Analysis of EFA6 distribution in axons and dendrites

E18 Cortical neurons were fixed at DIV 3, 7, 14 or 21 and EFA6 was detected using the antibody described above. The axon initial segment was located using anti-pan-neurofascin. All cultures were fixed and labelled using identical conditions. EFA6 fluorescence intensity was measured in the AIS and at a region >50μm beyond the AIS, and then at similar regions in two dendrites to give a mean dendrite figure. Images were acquired by confocal laser scanning microscopy using a Leica TCS SPE confocal microscope. Identical settings were used for the acquisition of each image using Leica LAS AF software. Z-stacks were acquired for each image, spanning the entire depth of each neuron. GraphPad Prism was used for statistical analysis of data using ANOVA followed by Bonferroni’s post-hoc analysis as indicated in the figure legends.

### Axonal ARF activation assay

Active ARF was detected using a peptide derived from the active ARF binding domain (ABD) of GGA3 fused to a GST tag (GGA3-ABD- GST, Thermo). Neurons were fixed for 15 minutes in 3% formaldehyde (TAAB) in PBS, permeabilised with 0.1% triton for two minutes and incubated with 20μg/ml GGA3-ABD- GST in TBS and 1mM EDTA overnight at 4°C. The GST tag was then detected using rabbit anti-GST (Abcam ab19256, 1:400) and standard immunofluorescence. Control and EFA6 shRNA treated cultures were fixed and labelled in parallel, using identical conditions. Axons were analysed 200-1000μm distal to the cell body. Images of axons were acquired by confocal laser scanning microscopy using a Leica TCS SPE confocal microscope. Initial observations were made and detection settings were adjusted so that the pixel intensities of acquired images were below saturation. Settings were then stored and were applied for the identical acquisition of each image using Leica LAS AF software. Z-stacks were acquired for each image, spanning the entire depth of each axon. Maximum projection images were created, and used for analysis. Lines were then traced along sections of axons to define the region of interest, and mean pixel intensities per axon section were quantified using Leica LAS AF software. The acquired images were corrected for background by subtracting an identical region of interest adjacent to the axon being analysed. The same technique was then used for measuring total levels of ARF6 in axons, after ARF6 immunolabelling. GraphPad Prism was used for statistical analysis of data using students’ T-test, as indicated in the figure legends.

### EFA6 shRNA validation

The efficacy of shRNA targeting EFA6 (target sequence cagtcctggattactcgcatcaatgtggt) was confirmed by immunofluorescence and western blotting. E18 cortical neurons were transfected with shRNA targeting EFA6 or non-effective scramble control shRNA at +10DIV, fixed at +14DIV and immunolabelled for EFA6. To confirm the silencing efficiency and target validity, we used an overexpression silencing and rescue approach similar to that described previously (Choi et al., 2006), using human EFA6 to rescue knockdown as there are base pair differences in the equivalent human sequence (cagtcctggat<u>C</u>actcgcatcaatgt<u>A</u>gt) compared to the rat. Preliminary experiments found that shRNA targeting rat EFA6 had no effect on human EFA6 expression levels in PC12 cells stably expressing human EFA6. We did not use this to rescue silencing in primary neurons because we found that overexpressed EFA6 localised erroneously throughout the axon. To confirm silencing by western blotting, PC12 cells were transfected with either rat EFA6-FLAG or rat EFA6-FLAG plus human EFA6-FLAG (as an shRNA resistant rescue plasmid) together with control or EFA6 shRNA, and lysates were used for western blotting (human and rat EFA6 run at the same size on western blots (Choi et al., 2006)). Expression levels of EFA6 were determined by immunoblotting with anti-FLAG. RFP and actin were probed for normalisation.

### Analysis of microtubule dynamics using EB3-GFP

Cortical neurons were transfected at DIV10 with EB3-GFP, and imaged at DIV 13-15 using spinning disc confocal microscopy. Images were acquired of cell bodies and initial sections of axons, and subsequently of regions of axons more distal into the axon (200-400μm). Images were acquired every second for three minutes. Kymographs were generated using MetaMorph software, and used to quantify the dynamics of EB3 comets. Velocity, duration, length and number of comets were measured per axon section. ANOVA analysis confirmed there was no variation in the length of axon sections analysed. GraphPad Prism was used for statistical analysis of data using ANOVA and Bonferroni’s post-hoc analysis.

### Live imaging of α9 integrin, rab11 and APP for axon transport analysis

Cortical neurons were transfected at DIV10 with either α9 integrin-GFP, rab11-GFP or APP-GFP together with either control or EFA6 shRNA and imaged at DIV 14-15 using spinning disc confocal microscopy. For α9 integrin-GFP and rab11-GFP, sections of axons were imaged at the AIS, at a region in the proximal part of the axon (100-300μm) and a region in the distal part of the axon (>600μM). APP-GFP expressing axons were imaged at a region in the proximal part of the axon (100-300μm) and a region in the distal part of the axon (>600μM). Vesicles were tracked for their visible lifetime, and analyzed by kymography to determine the amount of vesicles classed as anterograde, retrograde, bidirectional or immobile per axon section. This was determined as described previously (Eva et al., 2012; Franssen et al., 2015): vesicles with a total movement less than 2 μm during their visible lifetimes were classed as immobile. Vesicles moving in both directions but with net movement of less than 2 μm (during their visible lifetimes) were classed as bidirectional, even though total movement may have been larger. Vesicles with net movements greater than 5 μm in either direction by the end of their visible lifetimes were classed as anterograde or retrograde accordingly. ANOVA analysis confirmed there was no difference in the length of axon sections analysed. GraphPad Prism was used for statistical analysis of data using ANOVA and Bonferroni’s post-hoc analysis.

### Measurement of β1 integrin and rab11 axon-dendrite ratio

Cortical neurons were transfected at DIV10 with either control or EFA6 shRNA and fixed at DIV14. These were immunolabelled for either β1 integrin or rab11. Control and EFA6 shRNA treated cultures were fixed and labelled in parallel, using identical conditions. Images were acquired by confocal laser scanning microscopy using a Leica TCS SPE confocal microscope, using identical settings for each image (determined separately for either β1 integrin or rab11). Images were acquired at 40x to include the cell body, dendrites and a section of axon in each image. Leica LAS AF software was used to measure mean fluorescnece in the axon, and in two dendrites (to give a mean dendrite measurement). A region next to each neurite was used to subtract background fluorescence. Axon dendrite-ratio was determined as the mean dendrite fluorescence intensity divided by the axon intensity. GraphPad Prism was used for statistical analysis of data usinga students T-Test.

### Axonal identification

Axons and/or axon initial segments were identified either by immunolabelling of cultures with fluorescently conjugated anti-pan-neurofascin (Franssen et al., 2015), or by morphology (axons of transfected cells were clearly distinguishable from dendrites due to their length and lack of spines at more mature stages). For live labelling of neurofascin, neurons were live labelled with an Alexa-conjugated antibody (488, 594 or 350nm) for 45 min at 37°C. For antibody conjugation, the Tris-containing buffer of anti-neurofascin was first exchanged into 0.05M borate buffer by dialysis using a D-tube dialyzer (Novagen) and the antibody was then fluorescently labelled using a DyLight Antibody Labeling Kit (Thermo).

### Laser Axotomy of Cultured Neurons

Axons were severed in vitro using a 355 nm DPSL laser (Rapp OptoElectronic, Hamburg, Germany) connected to a Leica DMI6000B microscope. Cortical neurons were axotomised at DIV14-17 at distance of 800-200μm distal to the cell body on a section of axon free from branches. In this way axons were cut at a substantial distance from the cell body, but not close to the end of the axon. Only highly polarised neurons (with many dendrites and a single axon) were chosen. A single axon cut was made per neuron. Images after axotomy were acquired every 30 minutes for 14 hours. The response to axotomy was recorded as regeneration or fail. Regeneration was classed as the development of a new growth cone followed by axon extension for a minimum of 50μm. Fail was then classed as either bulb or stump phenotype. Regenerated axons were analysed for the time taken to regeneration, growth cone area and length of axon growth in the 14 hours following axotomy. Data regarding regeneration percentage of cortical axons was analysed by Fisher’s exact test. GraphPad Prism was used for statistical analysis of the remaining data. Adult DRG neurons were axotomised as described above, except that the location for axotomy was chosen as directly before a growth cone, so as to determine the proportion of axons that would regenerate a growth cone rapidly after injury. Data were analysed using Fisher’s exact test.

### Statistical Analysis

Statistical analysis was performed throughout using Graphpad Prism. Fisher’s exact test was calculated using Graphpad online: https://www.graphpad.com/quickcalcs/contingency1.cfm Data were analysed using ANOVA with post hoc analysis, Students T-Test, and Fisher’s exact test. For analysis of EFA6 distribution in axons and dendrites, the axon and two dendrites from 15 neurons were analysed, from three separate sets of primary cultures: total 45 neurons analysed. Statistical analysis was performed using ANOVA followed by Tukey's multiple comparison test as indicated in the figure legends. For quantification of axonal ARF activity in cortical neurons, 61 vs 64 axons were analysed, from three separate experiments. 50 axons were analysed for total axonal ARF6 quantification, from three experiments. For cortical vs. DRG axon ARF quantification, n=58 (DRG) vs. 60 (cortical), from three separate experiments. Data were analysed by student’s t-test. For analysis of EB3-GFP comets, 30 vs. 40 axon sections were imaged and analysed. Data were analysed by student’s t-test. Quantification of α9 integrin-GFP axon transport: n=12 to24 neurons per condition, from more than three experiments, analysed using Anova and Bonferroni’s comparison test. Quantification of β1integrin axon-dendrite ratio: n=72 neurons in total from three experiments, analysed by student’s t-test. Quantification of rab11-GFP axon transport: n=19 to27 neurons per condition from more than three experiments. Quantification of rab11 axon-dendrite ratio: n=53 vs. 55 neurons from three experiments, analysed by students t-test. Cortical neuron axon regeneration analysis: n=59 vs. 63 neurons from more than three experiments, percentage regenerating was compared by fisher’s exact test. Percentage of failed axons bulb vs. stump, was compared by fisher’s exact test. Distance grown after regeneration, area of regenerated growth cones, and time taken to establish a growth cone, were each compared by student’s t-test. Quantification of axon regeneration of DRG neurons: n=48 expressing GFP, n=44 expressing EFA6-GFP or n=31 expressing EFA6 E242K. Data were analysed by fisher’s exact test.

## Acknowledgements

Acknowledgements: The study was funded by grants from the Christopher and Dana Reeve Foundation, the Medical Research Council, the ERC advanced grant ECMneuro, the International Spinal Research Trust, Glaxo Smith Kline International Scholarship, Honjo International Scholarship, Bristol-Myers Squibb Graduate Studentship and the NIHR Cambridge Biomedical Research Centre. We thank Eunjoon Kim (Daejeon) for EFA6 antibodies, Casper Hoogenraad (Utrecht) for EB3-GFP, and Frederic Luton (Valbonne) for EFA6 E242K-GFP.

## Supplementary Figures and Legends

**Fig. S1.**
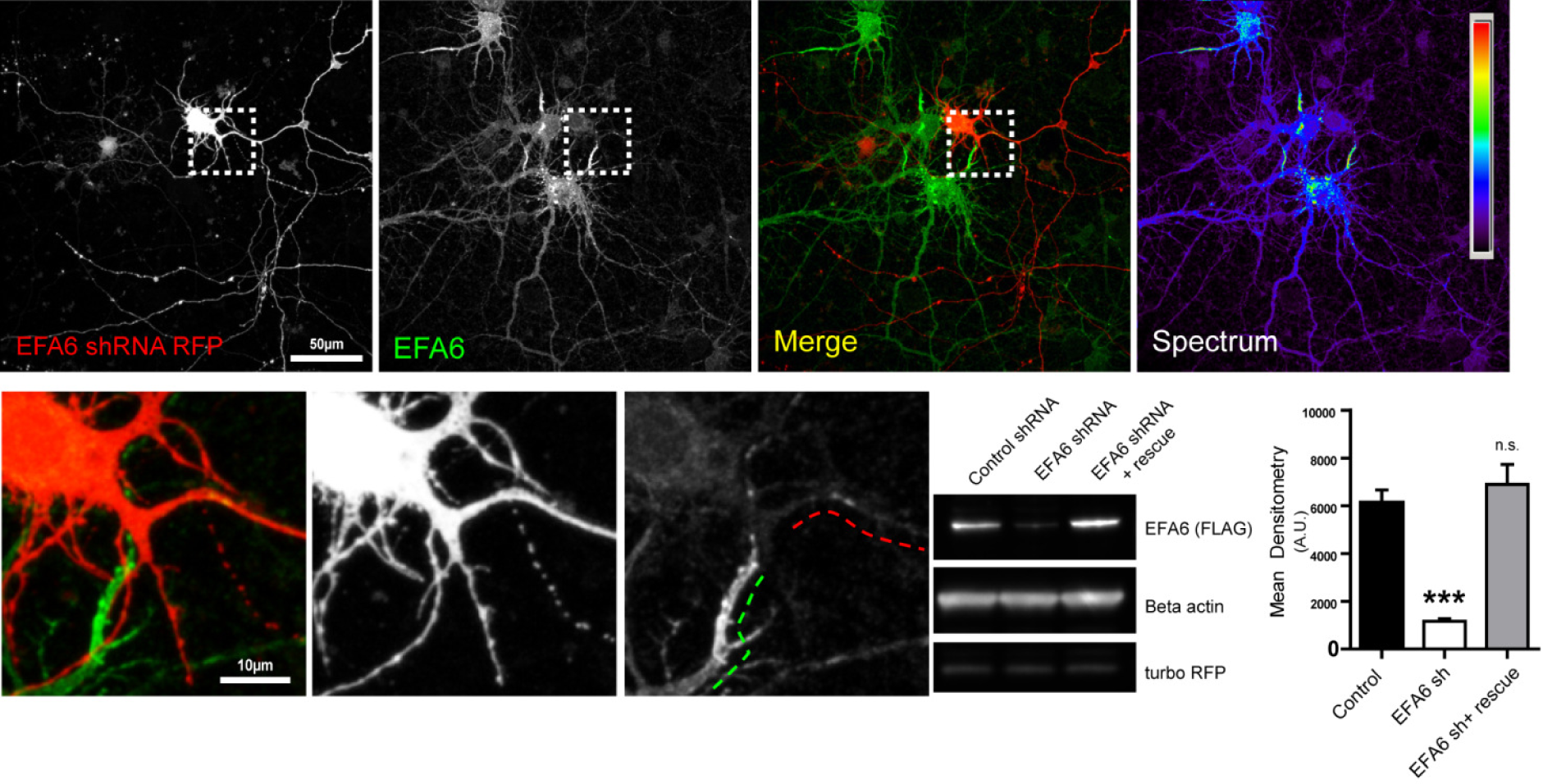
Validation of shRNA targeting EFA6. E18 cortical neurons transfected with shRNA targeting EFA6 (red, RFP) at +10DIV, fixed at +14DIV and immunolabelled for EFA6. Lower panels are high magnification of the region indicated by the dashed box in the upper panels, showing low levels of EFA6 in the initial part of the axon of a transfected cell (red dashes), compared with high levels of EFA6 in the untransfected cell (dashed green lines). **(B)** Validation of EFA6 shRNA by western blotting. Blots show lysate from PC12 cells transfected with either rat EFA6-FLAG and or rat EFA6- FLAG plus human EFA6- FLAG (as an shRNA resistant rescue plasmid) together with control or EFA6 shRNA. Graph is quantification of silencing effect by densitometry. ***p=0.0002, f=25.8, ANOVA and Bonferroni’s test.

**Fig. S2.**
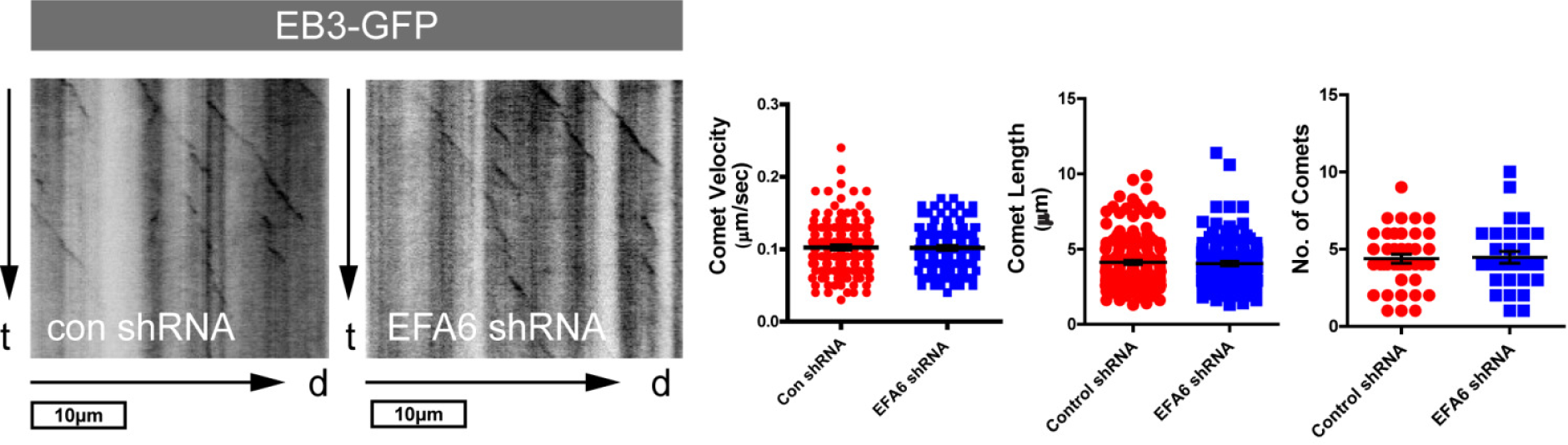
EFA6 silencing does not alter axon microtubule dynamics. Representative kymographs and analysis of EB3-GFP comets in axons of E18+14DIV cortical neurons co-expressing EB3-GFP and control or EFA6 shRNA. n = 30 to 40 neurons. EB3-GFP localised strongly to the AIS, however comets were undetectable here in cells expressing either control or EFA6 shRNA (see movies S1 and S2).

**Fig. S3.**
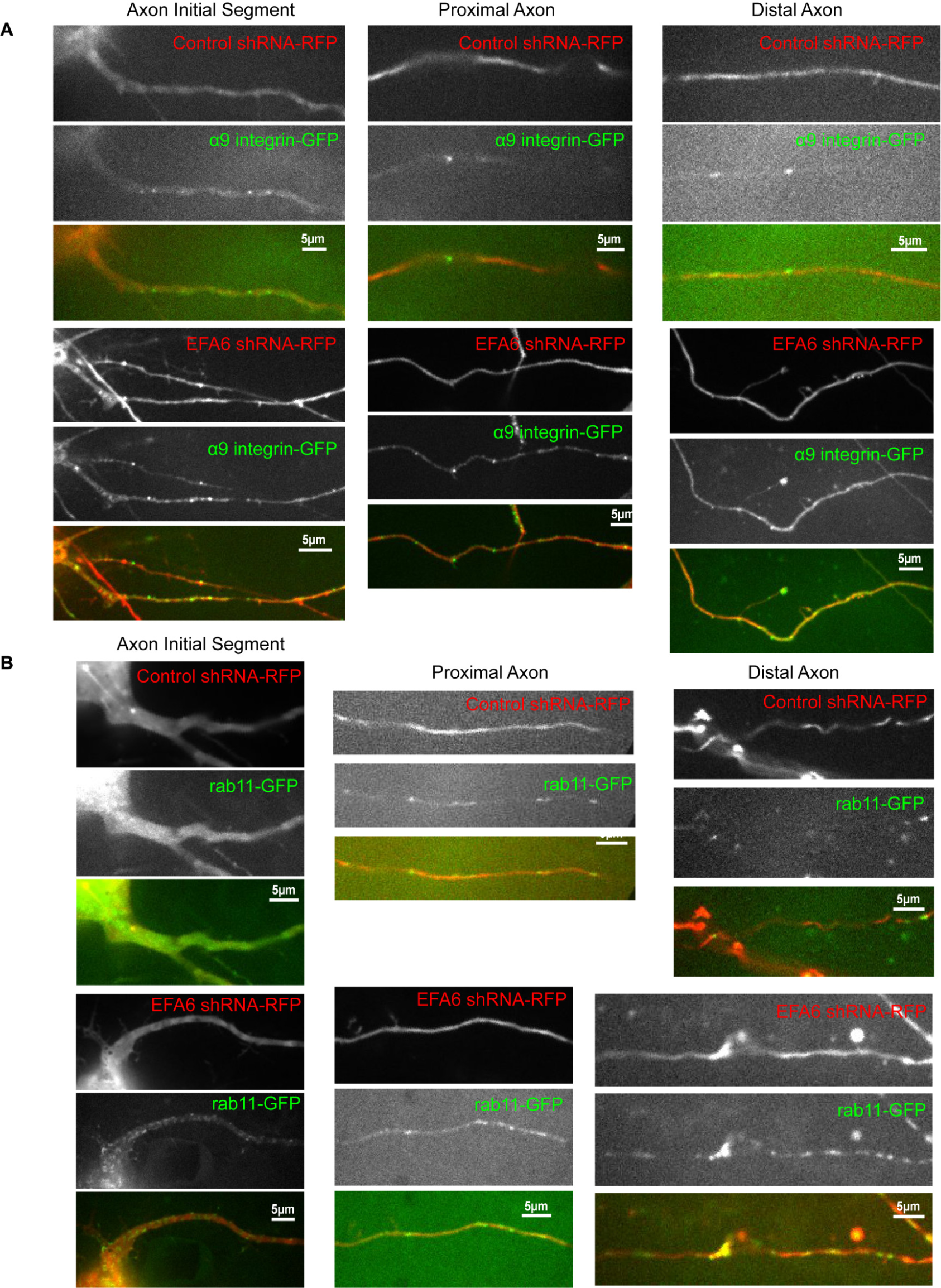
Single frame images of movies used to analyse axon endosome dynamics. Images show sections of the initial proximal or distal part of axons, expressing α9 integrin-GFP or rab11-GFP together with either control or EFA6 shRNA. **(A)** Axons of E18 +14-17DIV cortical neurons transfected at +10DIV with control or EFA6-shRNA, and co-transfected with α9 integrin-GFP. **(B)** Axons of E18 +14-17DIV cortical neurons transfected at +10DIV with control or EFA6-shRNA, and co-transfected with rab11-GFP.

**Fig. S4.**
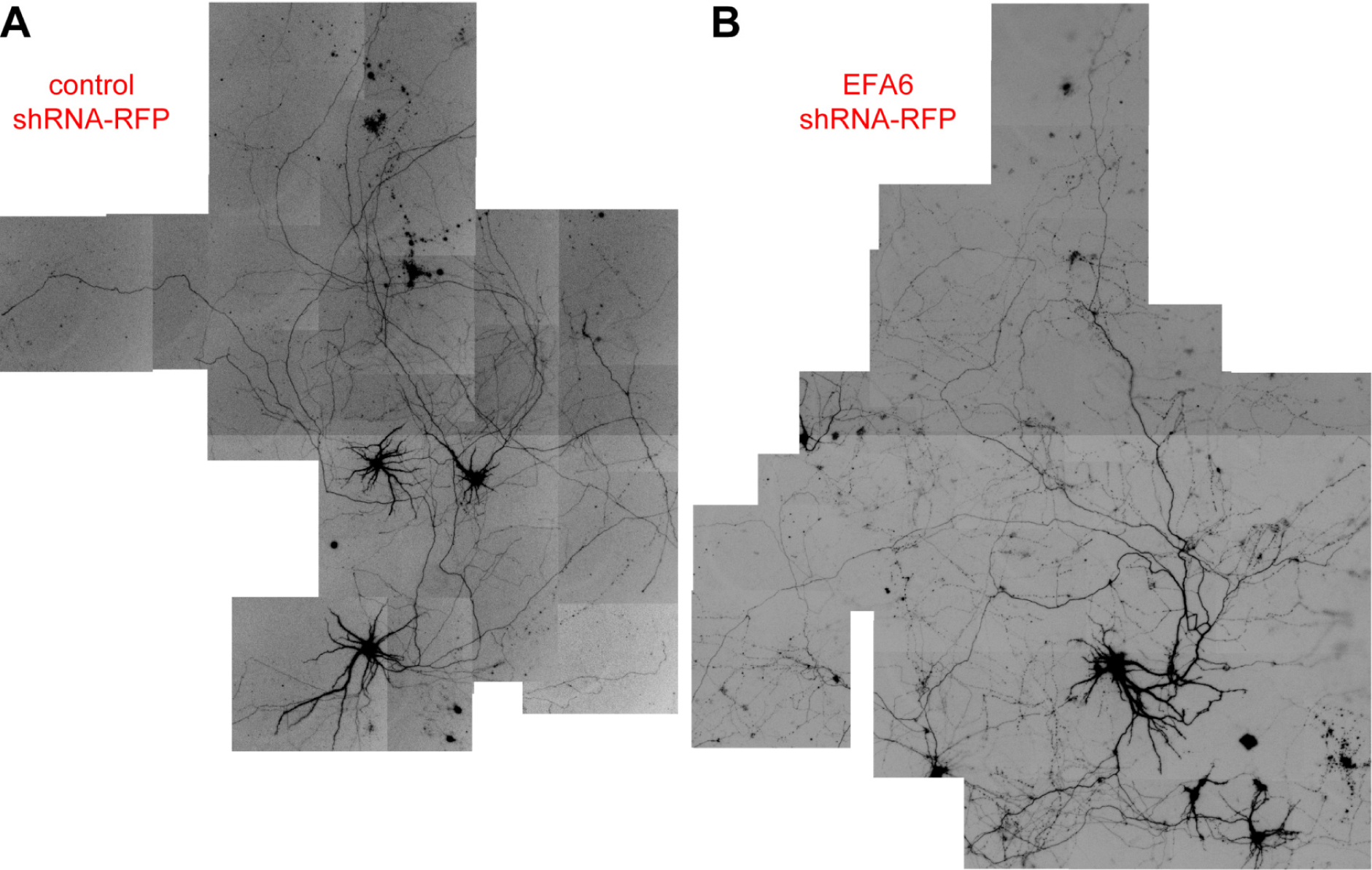
Examples of control- or EFA6-shRNA expressing neurons used for axotomy experiments. **(A)** Example of cortical neurons transfected with control shRNA-RFP (signal is RFP) at E18 +10DIV, used for axotomy experiments at E18 +14-17DIV. **(B)** Example of cortical neurons transfected with EFA6 shRNA-RFP (signal is RFP) at E18 +10DIV, used for axotomy experiments at E18 +14-17DIV.

**Fig. S5.**
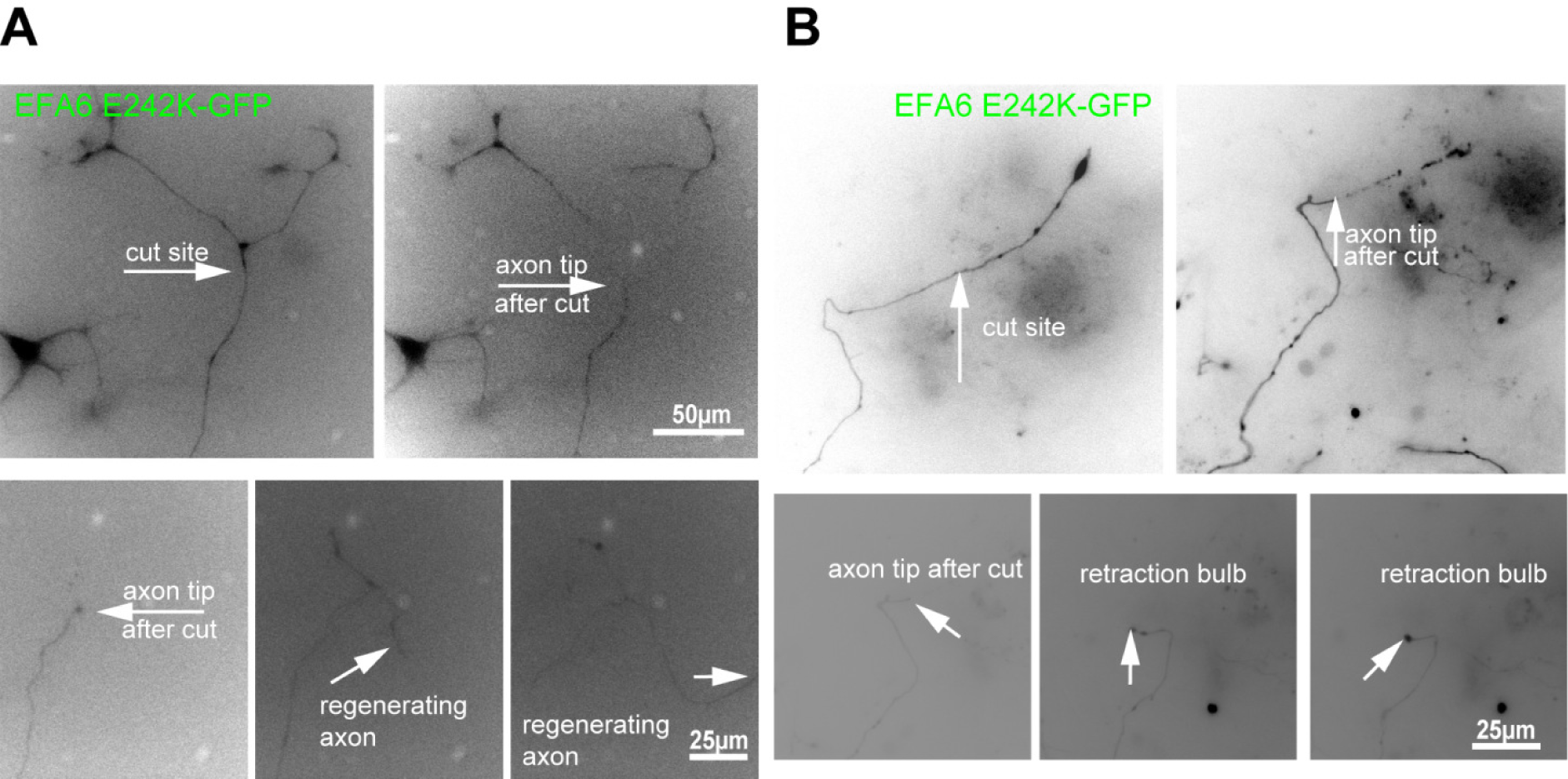
DRG neurons expressing EFA6 with inactive GEF domain (EFA6 E242K). **(A)** Example of successful regeneration after axotomy, DRG neuron expressing EFA6 E242K. **(B)** Example of regeneration failure after axotomy, DRG neuron expressing EFA6 E242K. Expression of EFA6 E242K allows 50.5% of axons to regenerate their growth cones after axotomy.

## Movie Legends

**Movie 1.** EB3-GFP localised to the axon initial segment and cell body (comets) of E18 +15DIV cortical neuron, also expressing control shRNA.

**Movie 2.** EB3-GFP localised to the axon initial segment and cell body (comets) of E18 +15DIV cortical neuron, also expressing shRNA targeting EFA6. The apparent break in the axon is a section of axon out of the plane of focus.

**Movie 3.** α9 integrin axon transport in the distal section of an axon also expressing control shRNA (see fig. S2). Movement to the left hand side is retrograde, right hand side anterograde.

**Movie 4.** α9 integrin axon transport in the distal section of an axon also expressing shRNA targeting EFA6 (see fig. S2). Movement to the left hand side is retrograde, right hand side anterograde.

**Movie 5.** Axon regeneration after laser axotomy of E18 +15DIV cortical neuron expressing control shRNA.

**Movie 6.** Axon regeneration after laser axotomy of E18 +15DIV cortical neuron expressing shRNA targeting EFA6.

**Movie 7.** Axon regeneration after laser axotomy of adult DRG neuron expressing GFP.

**Movie 8.** Failed axon regeneration after laser axotomy of adult DRG neuron overexpressing EFA6-GFP.

